# A phylogenomic framework and timescale for comparative studies of tunicates

**DOI:** 10.1101/236448

**Authors:** Frédéric Delsuc, Hervé Philippe, Georgia Tsagkogeorga, Paul Simion, Marie-Ka Tilak, Xavier Turon, Susanna López-Legentil, Jacques Piette, Patrick Lemaire, Emmanuel J. P. Douzery

**Affiliations:** ISEM, Université de Montpellier, CNRS, IRD, EPHE, Montpellier, France; Centre for Biodiversity Theory and Modelling, UMR CNRS 5321, Station d’Ecologie Théorique et Expérimentale, Moulis, France; Département de Biochimie, Centre Robert-Cedergren, Université de Montréal, Montréal, Canada; School of Biological and Chemical Sciences, Queen Mary University of London, London, UK; Center for Advanced Studies of Blanes (CEAB, CSIC), Girona, Spain; Department of Biology and Marine Biology, Center for Marine Science, University of North Carolina Wilmington, Wilmington, NC, USA; Centre de Recherche en Biologie cellulaire de Montpellier, UMR 5237, CNRS, Université de Montpellier, Montpellier, France

**Keywords:** Tunicata, Thaliacea, Molecular dating, Transcriptomes, Phylogenomics, Evo-devo

## Abstract

**Background:** Tunicates are the closest relatives of vertebrates and are widely used as models to study the evolutionary developmental biology of chordates. Their phylogeny, however, remains poorly understood and to date, only the 18S rRNA nuclear gene and mitogenomes have been used to delineate the major groups of tunicates. To resolve their evolutionary relationships and provide a first estimate of their divergence times, we used a transcriptomic approach to build a phylogenomic dataset including all major tunicate lineages, consisting of 258 evolutionarily conserved orthologous genes from representative species.

**Results:** Phylogenetic analyses using site-heterogeneous CAT mixture models of amino acid sequence evolution resulted in a strongly supported tree topology resolving the relationships among four major tunicate clades: 1) Appendicularia, 2) Thaliacea + Phlebobranchia + Aplousobranchia, 3) Molgulidae, and 4) Styelidae + Pyuridae. Notably, the morphologically derived Thaliacea are confirmed as the sister-group of the clade uniting Phlebobranchia + Aplousobranchia within which the precise position of the model ascidian genus *Ciona* remains uncertain. Relaxed molecular clock analyses accommodating the accelerated evolutionary rate of tunicates reveal ancient diversification (~450-350 million years ago) among the major groups and allow comparing their evolutionary age with respect to the major vertebrate model lineages.

**Conclusions:** Our study represents the most comprehensive phylogenomic dataset for the main tunicate lineages. It offers a reference phylogenetic framework and first tentative timescale for tunicates, allowing the direct comparison with vertebrate model species in comparative genomics and evolutionary developmental biology studies.

## Background

Large-scale phylogenetic analyses of tunicate genomic data from a handful of model species have identified this marine chordate group as the closest relative of vertebrates [1–5]. This discovery has had profound implications for comparative genomics and evolutionary developmental biology studies aimed at understanding the origins of chordates and vertebrates [6–8]. Indeed, the new chordate phylogeny implies that the tunicate body plan is evolutionarily derived and has become secondarily simplified from more complex chordate ancestors [2,3].

The key phylogenetic position of tunicates within chordates has prompted the selection of model species such as *Ciona robusta* (formerly *Ciona intestinalis* type A [9]) for which a full genome has been sequenced early in the history of comparative genomics to provide insight into vertebrate-specific whole genome duplications [10]. Since then, genome sequences have been assembled for additional species that are widely used as models in comparative genomics and evolutionary developmental biology [11] including *Ciona savignyi* [12], *Oikopleura dioica* [5], *Botryllus schlosseri* [13], *Molgula occidentalis*, *M. occulta* and *M. occulata* [14], *Phallusia mammillata* [15] and *Halocynthia roretzi* [15]. The available genomic data have notably revealed a stunning contrast in the evolutionary rate of nuclear protein-coding genes between tunicates and vertebrates [3,16]. This accelerated evolution of tunicate genes is also coupled with extensive structural rearrangements observed in their genomes [5,17,18]. This contrast is even more pronounced for mitochondrial genomes, which are particularly fast evolving and highly rearranged in tunicates with respect to other deuterostomes in which they are widely conserved [5,19,20]. The reasons behind the rapid rate of genomic evolution in tunicates remain unclear [16,21,22] and contrast with the unusual conservation level of embryonic morphologies between all ascidian species studied so far [7].

Despite renewed interest in tunicate evolution, phylogenetic relationships among the major tunicate lineages remain uncertain. Previous molecular phylogenetic studies relying on 18S rRNA [23–26] and mitogenomes [20,27,28] have proposed first delineations of major tunicate clades, revoking the traditional 19^th^ century classification into the three classes Appendicularia (larvaceans), Thaliacea (salps, doliolids, and pyrosomes), and Ascidiacea (phlebobranchs, aplousobranchs, and stolidobranchs). Indeed, these studies found unanimous support for the paraphyly of Ascidiacea (ascidians) owing to the inclusion of thaliaceans in a clade also containing two main ascidian lineages (phlebobranchs and aplousobranchs) to the exclusion of stolidobranch ascidians (molgulids, pyurids, and styelids). Nevertheless, the resolving power of these standard markers — nuclear ribosomal RNA and mitochondrial protein-coding genes — appeared to be limited regarding the relationships among the three newly proposed main clades: 1) Appendicularia, 2) Stolidobranchia, and 3) Phlebobranchia + Thaliacea + Aplousobranchia. Notably, the relationships within the latter group were left unresolved, with the position of thaliaceans relative to phlebobranchs and aplousobranchs still being debated [25,27].

The phylogenetic position of thaliaceans is key for understanding the evolution of developmental modes within tunicates [29]. Compared to their closest relatives that are mostly solitary and sessile, the three groups of thaliaceans (salps, doliolids and pyrosomes) are pelagic with complex life cycles including solitary and colonial phases. Their unique lifestyle also seems to be associated with spectacular differences in their embryology, such as the loss of a well-developed notochord in the larva of most thaliaceans, with the exception of only a few doliolid species [29]. Based on our current understanding of tunicate evolution, thaliaceans may have evolved from a sessile ascidian-like ancestor and therefore can serve as a model to understand how the transition from a benthic to a pelagic lifestyle has led to drastic modifications in the morphology, embryology, and life cycle of these tunicates [29]. Coloniality is another remarkable feature of the thaliaceans, which shows some similarities with the coloniality in ascidians, even tough this trait probably evolved independently in the two groups [29]. It is noteworthy that doliolids have polymorphic colonies [30], a trait that is absent in colonial ascidians. A reliable phylogeny positioning thaliaceans with regard to colonial ascidians is thus necessary to understand the evolution of these unique features.

Outstanding questions in chordate evolution include the identification of the determinants of the rapid rate of genome evolution in tunicates and the emergence of vertebrates [11,31]. A prerequisite to address these issues is to reconstruct a reliable phylogenetic framework and timescale to guide future comparative evolutionary genomic and evolutionary studies of chordate development. Moreover, given that the fossil record of tunicates is deceptively scarce and controversial [32–34], a molecular timescale for chordates would allow comparing tunicate evolution to the well-calibrated vertebrates [35] for the first time. A phylogenetic and timing framework is notably critical for the identification and interpretation of both conserved and divergent developmental features of tunicates compared to model vertebrate species in the context of their fast rate of genomic evolution [11].

Here, we use new transcriptomic data obtained through high-throughput sequencing technologies (Roche 454 and Illumina HiSeq) to build the first tunicate phylogenomic dataset including all major tunicate groups. This dataset consists of 258 orthologous nuclear genes for 63 taxa including representative deuterostome species and all major chordate lineages. Using phylogenetic analyses based on the best-fitting site-heterogeneous CAT mixture model of amino acid sequence evolution, we inferred well-resolved phylogenetic relationships for the major clades of tunicates. Our molecular dating analyses based on models of clock relaxation accounting for variation in lineage-specific evolutionary rates provide a first tentative timescale for the emergence of the main tunicate clades allowing a direct comparison with vertebrate model systems.

## Methods

### Transcriptome data collection

Live tunicate specimens were ordered from Gulf Specimen Marine Laboratories, Inc. (Panacea, FL, USA) and the Roscoff Biological Station (France) services, and collected in Villefranche-sur-Mer (France), and Blanes (Spain). One single run of Roche 454 GS-FLX Titanium was conducted at GATC Biotech (Konstanz, Germany) on multiplexed total RNA libraries that were constructed for *Clavelina lepadiformis, Cystodytes dellechiajei, Bostrichobranchus pilularis, Molgula manhattensis, Molgula occidentalis, Phallusia mammillata, Dendrodoa grossularia, Polyandrocarpa anguinea,* and *Styela plicata*. Complementary RNAseq data were acquired with paired-end 100-nt Illumina reads at Beijing Genome Institute (Schenzen, China) for the thaliaceans *Salpa fusiformis* (mix of 2 blastozooids) and *Doliolum nationalis* (mix of 15 phorozooids), and with single-end 100-nt Illumina reads at GATC Biotech (Konstanz, Germany) for *Clavelina lepadiformis* and *Cystodytes dellechiajei* (mix of several individuals) [36]. Previously obtained 454 transcriptomic data for *Microcosmus squamiger* [16] were also considered. De novo assemblies were conducted with Trinity [37] for 454 reads, and ABySS [38] for Illumina reads using the programs’ default parameters. For both kinds of libraries, we confirmed the sample taxonomic identifications by assembling the mitochondrial CO1 and nuclear 18S rRNA barcoding genes and reconstructing maximum likelihood trees with available comparative data. Additional tunicate sequences were collected in public databases from various sequencing projects: *Botryllus schlosseri*, *Halocynthia roretzi*, and *Diplosoma listerianum* (ESTs), *Molgula tectiformis* (cDNAs), and *Ciona robusta*, *Ciona savignyi*, and *Oikopleura dioica* (genomes). Detailed information on biological specimens, basic statistics, and accession numbers of newly sequenced transcriptomes can be found in Additional file 1: Table S1.

### Phylogenomic dataset assembly

We built upon a previous phylogenomic dataset [39] to select a curated set of 258 orthologous markers for deuterostomes. Alignments were complemented with sequences from the NCBI databases using a multiple best reciprocal hit approach implemented in the newly designed Forty-Two software [40]. Because 454 DNA sequence reads are characterized by sequencing errors typically disrupting the reading frame when translated into amino acids, alignments were verified by eye using the program ED from the MUST package [41]. Ambiguously aligned regions were excluded for each individual protein using Gblocks with medium default parameters [42] with a few subsequent manual refinements using NET from the MUST package to relax the fact that this automated approach is sometimes too conservative. This manual refinement step restored only 418 amino acid sites (i.e., 0.6 % of the total alignment length). Potential cross-contaminations between our samples were also dealt with at the alignment stage by performing BLAST searches of each sequence against a taxon-rich reference database maintained for each curated gene alignment, and were further looked for by a visual examination of each individual gene phylogeny.

The concatenation of the resulting 258 amino acid alignments was constructed with ScaFos [43] by defining 63 deuterostomian operational taxonomic units (OTUs) representing all major lineages. The taxon sampling included 18 tunicates, 34 vertebrates, one cephalochordate, with seven echinoderms, two hemichordates and one xenoturbellid as more distant outgroups. When several sequences were available for a given OTU, the slowest evolving one was selected by ScaFos, according to ML distances computed by TREE-PUZZLE [44] under a WAG+F model. The percentage of missing data per taxon was reduced by creating some chimerical sequences from closely related species (i.e., *Eptatretus burgeri* / *Myxine glutinosa*, *Petromyzon marinus* / *Lethenteron japonicum*, *Callorhinchus milii* / *C. callorynchus*, *Latimeria menadoensis* / *L. chalumnae*, *Rana chensinensis* / *R. catesbeiana*, *Alligator sinensis* / *A. mississippiensis*, *Chrysemys picta* / *Emys orbicularis* / *Trachemys scripta, Patiria miniata* / *P. pectinifera* / *Solaster stimpsonii*, *Apostichopus japonicus* / *Parastichopus parvimensis*, *Ophionotus victoriae* / *Amphiura filiformis*) and by retaining only proteins with at most 15 missing OTUs. The tunicate *Microcosmus squamiger* was excluded at this stage due to a high percentage of missing data resulting from the low number of contigs obtained in the assembly. The final alignment comprised 258 proteins and 63 taxa for 66,593 unambiguously aligned amino-acid sites with 20% missing amino acid data.

### Phylogenetic analyses

Bayesian cross-validation [45] implemented in PhyloBayes 3.3f [46] was used to compare the fit of site-homogeneous (LG and GTR) and site-heterogeneous (CAT-F81 and CAT-GTR) models coupled with a gamma distribution (Γ_4_) of site-rate heterogeneity. Ten replicates were considered, each one consisting of a random subsample of 10,000 sites for training the model and 2,000 sites for computing the cross-validation likelihood score. Under site-homogeneous LG+Γ_4_ and GTR+Γ_4_ models, 1,100 sampling cycles were run and a burn-in of 100 samples was used, and under site-heterogeneous models CAT-F81+Γ_4_ and CAT-GTR+Γ_4_, 3,100 sampling cycles were run and the first 2,100 samples were discarded as burn-in.

Bayesian phylogenetic reconstruction under the best-fitting CAT-GTR+Γ_4_ mixture model [47] was conducted using PhyloBayes_MPI 1.5a [48]. Two independent Markov Chain Monte Carlo (MCMC) starting from a randomly generated tree were run for 6,000 cycles with trees and associated model parameters being sampled every cycle. The initial 1,000 trees sampled in each MCMC run were discarded as burn-in after checking for convergence in both likelihood and model parameters, as well as in clade posterior probabilities using *bpcomp* (max_diff < 0.3). The 50% majority-rule Bayesian consensus tree and the associated posterior probabilities (PP) were then computed from the remaining combined 10,000 (2 × 5,000) trees using *bpcomp*.

We further assessed the robustness of our phylogenomic inference by applying a gene jackknife resampling procedure [3]. A hundred jackknife replicates constituted of 130 alignments drawn randomly out of the total 258 protein alignments were generated. The 100 resulting jackknife supermatrices were then analysed using PhyloBayes_MPI under the second best-fitting CAT-F81+Γ_4_ model instead of the best-fitting CAT-GTR+Γ_4_ and for 2,000 sampling cycles in order to reduce computational burden. After removing the first 200 sampled trees of each chain as the burn-in, a majority-rule consensus tree was obtained for each replicate using the 1,800 trees sampled from the posterior distribution. A consensus tree was then obtained from the 100 jackknife-resampled consensus trees. The support values displayed by this Bayesian consensus tree are thus gene Jackknife Support percentages (JS). High values indicate nodes that have high posterior probability support in most jackknife replicates and are thus robust to gene sampling. We verified convergence of MCMCs in each Jackknife replicate by checking that varying the burn-in value did not affect the JS percentages obtained in the final consensus.

### Molecular dating

Molecular dating analyses were performed in a Bayesian relaxed molecular clock framework using PhyloBayes 3.3f [46]. In all dating calculations, the tree topology was fixed to the majority-rule consensus tree inferred in previous Bayesian analyses (Fig. 1). Dating analyses were conducted using the best-fitting site-heterogeneous CAT-GTR+Γ_4_ mixture model and a relaxed clock model with a birth–death prior on divergence times combined with soft fossil calibrations following Lartillot et al. [46]. Given the lack of trustable fossils within tunicates, we used 12 calibration intervals defined within vertebrates [49,50] and one within echinoderms [51]: 1) Chordata (Max. Age: 581 Mya, Min. Age: 519 Mya); 2) Olfactores (Max: 581, Min: 519); 3) Vertebrata (Max: 581, Min: 461); 4) Gnathostomata (Max: 463 Mya, Min: 422); 5) Osteichthyes (Max: 422, Min: 416); 6) Tetrapoda (Max: 350, Min: 330); 7) Amniota (Max: 330, Min: 312); 8) Diapsida (Max: 300, Min: 256); 9) Batrachia (Max: 299, Min: 200); 10) Clupeocephala (Max: 165, Min: 150); 11) Mammalia (Max: 191, Min: 163); 12) Theria (Max: 171, Min: 124); and 13) Echinoidea (Min: 255). The prior on the root of the tree (Deuterostomia) was set to an exponential distribution of mean 540 Mya.

**Figure 1.**
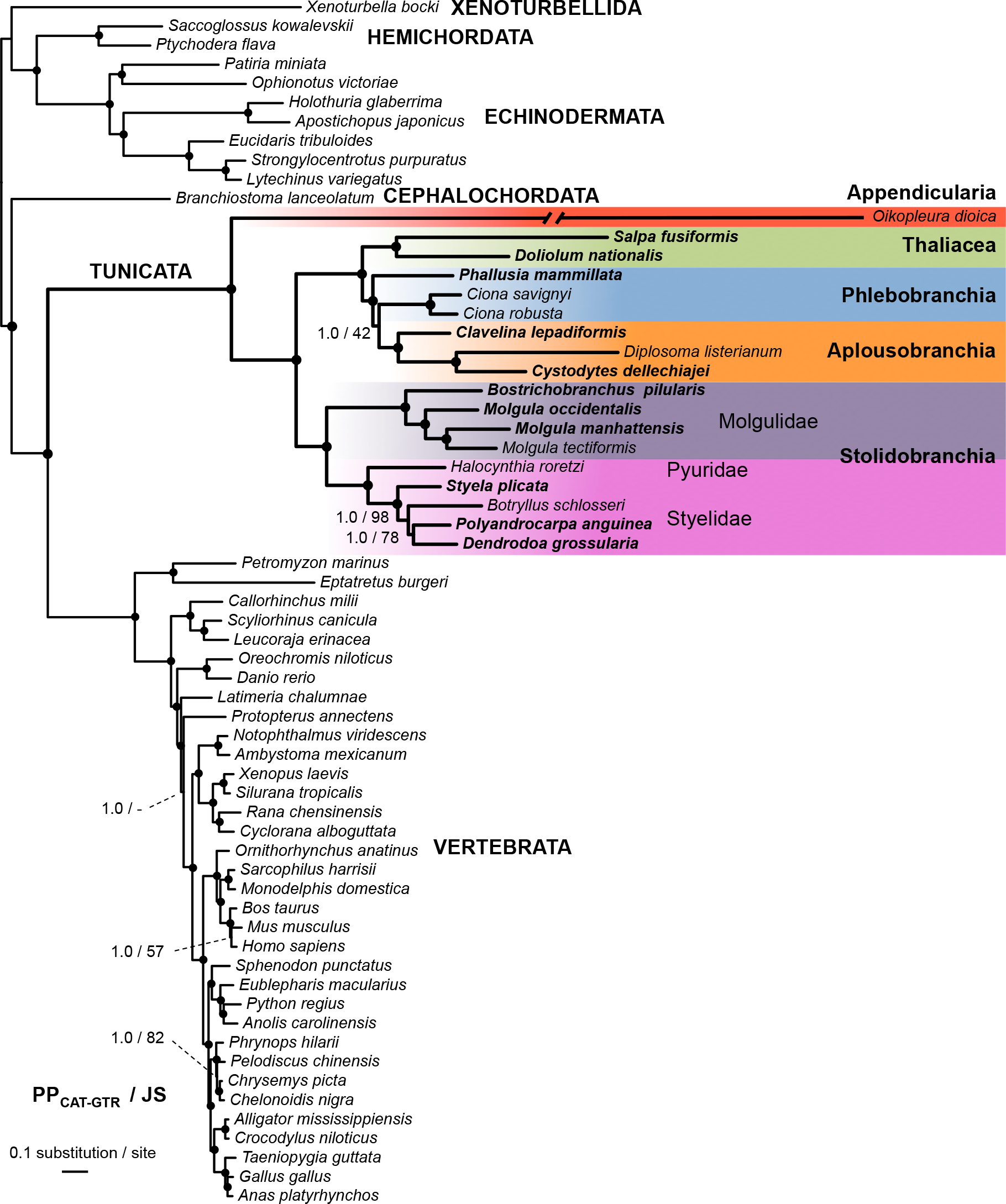
Phylogenetic relationships of 63 chordates highlighting the major tunicate groups inferred from 66,593 amino acid sites of 258 proteins. The Bayesian consensus phylogram has been inferred by PhyloBayes under the CAT-GTR+Γ_4_ mixture model. Values at nodes indicate Bayesian posterior probabilities (PP_CAT-GTR_) obtained under CAT-GTR+Γ_4_, and Jackknife Support percentages (JS), respectively. Circles at nodes pinpoint branches with maximal support from both methods. Species with newly obtained data are indicated in bold. The branch leading to the fast evolving *Oikopleura dioica* has been halved for graphical purposes.

In order to select the best-fitting clock model, we compared the auto-correlated log-normal (LN) relaxed clock model [52] with the uncorrelated gamma (UGAM) model [53], and a strict molecular clock (CL) model. These three clock models were compared against each other using the same prior settings (see above) in a cross-validation procedure as implemented in PhyloBayes following Lepage et al. [54]. However, to reduce the computational burden, the CAT-F81+Γ_4_ mixture model was used instead of CAT-GTR+Γ_4_. The cross-validation tests were performed by dividing the original alignment in two subsets with 90% of sites for the learning set (59,934 sites) and 10% of sites for the test set (6,659 sites). The overall procedure was repeated over 10 random splits for which a MCMC was run on the learning set for a total 4,000 cycles sampling posterior rates and dates every cycle.

The first 3,000 samples of each MCMC were excluded as the burn-in for calculating the cross-validation scores averaged across the 10 replicates.

The final dating calculations were conducted under both LN and UGAM relaxed-clock models and the CAT-GTR+Γ_4_ mixture model of sequence evolution by running MCMCs for a total 25,000 cycles sampling posterior rates and dates every 10 cycles. The first 500 samples of each MCMC were excluded as the burn-in after checking for convergence in both likelihood and model parameters using *readdiv*. Posterior estimates of divergence dates and associated 95% credibility intervals were then computed from the remaining 2,000 samples of each MCMC using *readdiv*. Additional dating calculations using the same sampling scheme were also conducted under the LN relaxed-clock model but using the less computationally intensive CAT-F81+Γ_4_ mixture model.

## Results and Discussion

### A reference phylogenetic framework for model tunicates

The evolutionary relationships of tunicates have long been a matter of debate. This is mainly because tunicates are characterized by an overall accelerated rate of evolution in their nuclear and mitochondrial genomes compared to other deuterostome species. This large lineage-specific variation in evolutionary rates among tunicates [16] could result in LBA artefacts, which hamper the reliable reconstruction of their phylogenetic relationships [55– 57]. Another contributing factor to our limited understanding of tunicate evolution is the uneven availability of genome data across different tunicate lineages. To address these limitations, we used (i) a wider taxon sampling encompassing all major tunicate lineages including two divergent thaliaceans, (ii) numerous nuclear genes to reduce stochastic error, and (iii) powerful site-heterogeneous models that generally offer the best fit to phylogenomic data and have the advantage to be least sensitive to LBA and other potential phylogenetic artefacts [39,58,59]. Accordingly, the results of our Bayesian cross-validation tests showed that the CAT-GTR+Γ_4_ mixture model offered the best statistical fit to the data (ΔlnL = 1,506 ± 98 compared to LG+Γ_4_), followed by the CAT-F81+Γ_4_ mixture model (ΔlnL = 817 ± 112 compared to LG+Γ_4_), and the GTR+Γ_4_ model (ΔlnL = 266 ± 41 compared to LG+Γ_4_).

The majority-rule consensus tree obtained using Bayesian phylogenetic reconstruction under the best-fitting CAT-GTR+Γ_4_ site-heterogeneous mixture model is thus presented in Figure 1. This well-supported phylogenetic tree has been rooted between Xenambulacraria (Xenoturbellida + Ambulacraria) and Chordata following the results of Philippe et al. [39] showing that Xenacoelomorpha (acoelomorphs + xenoturbellids) were related to Ambulacraria (hemichordates + echinoderms) within Deuterostomia. These results have been recently challenged by two studies claiming support for a more external position of Xenacoelomorpha as a sister-group to Nephrozoa (Protostomia + Deuterostomia) [60,61]. However, this newly proposed position is still debated as it might be the result of a long-branch attraction (LBA) artefact caused by the very long branches of acoelomorphs in phylogenomic trees [62,63]. Hence, our choice to root our trees according to Philippe et al. [39], which in any case does not affect the phylogenetic relationships of chordates.

The inferred topology unambiguously recovered the monophyly of chordates (PP = 1.0; JS = 100) and grouped the reciprocally monophyletic tunicates and vertebrates into Olfactores to the exclusion of cephalochordates (PP = 1.0; JS = 100) in accordance with the newly established chordate phylogeny [1,3,4]. Within tunicates, the appendicularian *Oikopleura dioica* was the sister-group of all other included taxa (PP = 1.0; JS = 100). Within the latter, there was a well-supported split (PP = 1.0; JS = 100) between Stolidobranchia on one side, and Phlebobranchia, Aplousobranchia, and Thaliacea on the other side. The monophyletic Stolidobranchia included two main clades, the first corresponding to the family Molgulidae (PP = 1.0; JS = 100), and the second grouping the families Pyuridae and Styelidae (PP = 1.0; JS = 100). Within molgulids, *Bostrichobranchus pilularis* was the sister-group of the three species within the genus *Molgula* (PP = 1.0; JS = 100), while *M. occidentalis* was the sister-group of *M. manhattensis* + *M. tectiformis* (PP = 1.0; JS = 100). Lastly, the four styelids *Styela plicata*, *Botryllus schlosseri*, *Polyandrocarpa anguinea*, and *Dendrodoa grossularia* constituted a monophyletic group (PP = 1.0; JS = 100) with respect to the single species here representing pyurids (*Halocynthia roretzi*). Within styelids, *S. plicata* diverged first (PP = 1.0; JS = 98) followed by *B. schlosseri* as the sister-group of *P. anguinea* + *D. variolosus* (PP = 1.0; JS = 100). On the other side of the tree, Thaliacea branched with maximum statistical support (PP = 1.0; JS = 100) as the sister-group of the clade Phlebobranchia + Aplousobranchia. The traditional class-level taxon Ascidiacea — currently considered to embrace the orders Aplousobranchia, Phlebobranchia and Stolidobranchia [64] — therefore refers to a paraphyletic assemblage. An alternative classification scheme based on gonad position (not commonly used nowadays) recognized two orders within ascidians: Enterogona (corresponding to Phlebobranchia + Aplousobranchia) and Pleurogona (= Stolidobranchia) [30,65]. These alternative order-level taxa are recovered as monophyletic in our analyses. The three aplousobranchs analysed here unambiguously formed a monophyletic clade (PP = 1.0; JS = 100) with *Clavelina lepadiformis* being the sister-group of *Diplosoma listerianum* and *Cystodytes dellechiajei* (PP = 1.0; JS = 100). The phlebobranchs appeared as a paraphyletic group with the two *Ciona* species branching closer to the aplousobranchs than to the other phlebobranch species (*Phallusia mammillata*), although with no statistical support from the gene jackknife resampling analysis (PP = 100; JS = 42).

The results from this first phylogenomic study including all tunicate lineages were in line with recent studies [20,25–28] demonstrating that ascidians (Class Ascidiacea) form a paraphyletic group. Our results showed that phlebobranchs and aplousobranchs are undoubtedly closer to thaliaceans than to stolidobranchs (Fig. 1), and that a thorough taxonomic revision of the tunicate classes is necessary. It seems clear that the use of the Ascidiacea class should be abandoned in favour of more meaningful classification schemes. Even though the position of Thaliacea was not always statistically supported, it consistently appeared as the sister-group of phlebobranchs + aplousobranchs in previous studies [20,24– 26,28], except for a recent genome-scale study in which the positioning of *Salpa thompsoni* most likely suffered artefactual LBA attraction towards the fast-evolving appendicularians [66]. The robust phylogenetic position of thaliaceans found here indicates that they likely evolved from a sessile ancestor and their study can provide valuable information on the morphological transformations associated with the transition to the pelagic lifestyle [29].

The monophyly of the clade uniting phlebobranchs and aplousobranchs has never been challenged and thus we suggest to re-use the term Enterogona to define this group as originally proposed by Perrier [65] and subsequently redefined by Garstang [67]. The close relationship between thaliaceans and enterogones has also been supported by all previous molecular studies, as well as by morphological observations. The gonad position and the shared paired ontogenetic rudiment of the atrial cavity and opening might constitute two of their anatomical synapomorphies [68]. Lastly, we also confirmed the previously reported monophyly of stolidobranchs (= Pleurogona), with molgulids being the sister-group to styelids + pyurids.

Finally, our phylogenomic study casted new light on two recurring issues in tunicate phylogenetics. First, phlebobranchs have been repeatedly found to be paraphyletic, albeit usually with no statistical support [25–28,69], and the phylogenetic affinities among its members remains unclear. Notably, the traditional position of *Ciona* as a phlebobranch ascidian was challenged by Kott [70], who placed the genus within aplousobranchs on the basis of morphological characters. More recently, Turon & López-Legentil [69] and Shenkar et al. [28] found that *Ciona* was closer to aplousobranchs than to other phlebobranchs using mitochondrial DNA. These results are in agreement with the tree topology obtained in the present study, although it was not statistically supported. The positioning of the model *Ciona* genus and the phylogenetic relationships of phlebobranchs need to be the focus of additional phylogenomic studies including a denser taxon sampling. Second, although the position of appendicularians as sister-clade to all other tunicates was well supported here and in all previous tunicate phylogenomic studies [2,3], the extremely long branch of *Oikopleura dioica* coupled with our current inability to completely alleviate a potential LBA artefact — even with complex site-heterogeneous mixture models (see [59]) — prevent us from considering this species’ phylogenetic position as conclusive. The long appendicularian branch should be subdivided with the inclusion of additional divergent species in future phylogenomic analyses to definitively settle this point.

### Evolutionary rate variations and molecular clock models

As observed in previous phylogenomic studies of chordates [2–4], the Bayesian phylogram estimated under the best-fitting CAT-GTR+Γ_4_ mixture model revealed marked branch length heterogeneity (Fig. 1). The tunicate branch lengths not only were much longer than those of all the other deuterostome clades, but also displayed strong variations within tunicates. From the ancestral node of Olfactores, the tunicate median evolutionary rate as measured in terms of branch length was of 1.53 amino acid substitution per site compared to the vertebrate median evolutionary rate that was 0.65. From the ancestral vertebrate node, the average of branch lengths is 0.35 ± 0.05 amino acid replacements per site. In contrast, from the ancestral node of tunicates — excluding the super fast-evolving *Oikopleura dioica* — the average of branch lengths was 0.69 ± 0.19. For the proteins here combined for phylogenomic purposes, tunicates (to the exception of *Oikopleura dioica*) displayed a twice-higher rate of amino acid substitutions than vertebrates.

Such substitution rate variation among lineages — within tunicates, and between tunicates and other deuterostomes — needs to be accounted for in molecular dating analyses by using models of clock relaxation [52]. The selection of the clock model is often arbitrary and appears mostly dependent of the software choice, with an overwhelming majority of studies relying on the BEAST software [71] using an uncorrelated gamma (UGAM, also known as UCLN) model of clock relaxation. However, it has been shown that autocorrelated rate models, such as the autocorrelated log-normal model (LN), often provide a better fit with phylogenomic data [54,72,73]. Consequently, we compared the fit of both the UGAM and LN models to the fit of a strict molecular clock (CL) model for our dataset using cross-validation tests under the CAT-GTR+Γ_4_ model. As expected given the large lineage specific rate variation, both relaxed clock models largely outperformed the strict clock model (UGAM vs. CL: ΔlnL = 4,068 ± 125; LN vs. CL: ΔlnL = 4,057 ± 118). Among relaxed clock models, UGAM and LN were statistically equivalent in offering a very similar fit to our data (UGAM vs. LN: ΔlnL = 11 ± 38).

The use of a relaxed clock model allowed us to perform evolutionary rate comparisons in terms of number of substitutions per site per million years for the 63 terminal taxa considered (Fig. 2). The box plots clearly showed that tunicates evolved faster than other groups, especially compared to vertebrates that were the slowest evolving. On average, tunicates evolved 6.25 faster than vertebrates (two-tailed t test; *t* = 4.542, *p* = < 0.001*), 2.08 times faster than cephalochordates (two-tailed t test not applicable with only one cephalochordate), and 2.45 times faster than the outgroups (two-tailed t test; *t* = 1.711, *p* = 0.099^ns^) here included. The evolutionary rate variation was also much more pronounced within tunicates than within other groups, even when the very fast evolver *Oikopleura dioica* was excluded. For instance, the colonial species *Diplosoma listerianum* and *Salpa fusiformis* evolved considerably faster than the solitary species *Ciona* spp. and *Styela plicata*. This confirmed earlier observations based on a reduced number of taxa and substitution rate estimations on 35 housekeeping genes [16], once again underlining the peculiar genomic evolution of tunicates that might find its root in elevated mutation rates and pervasive molecular adaptation [21,22].

**Figure 2.**
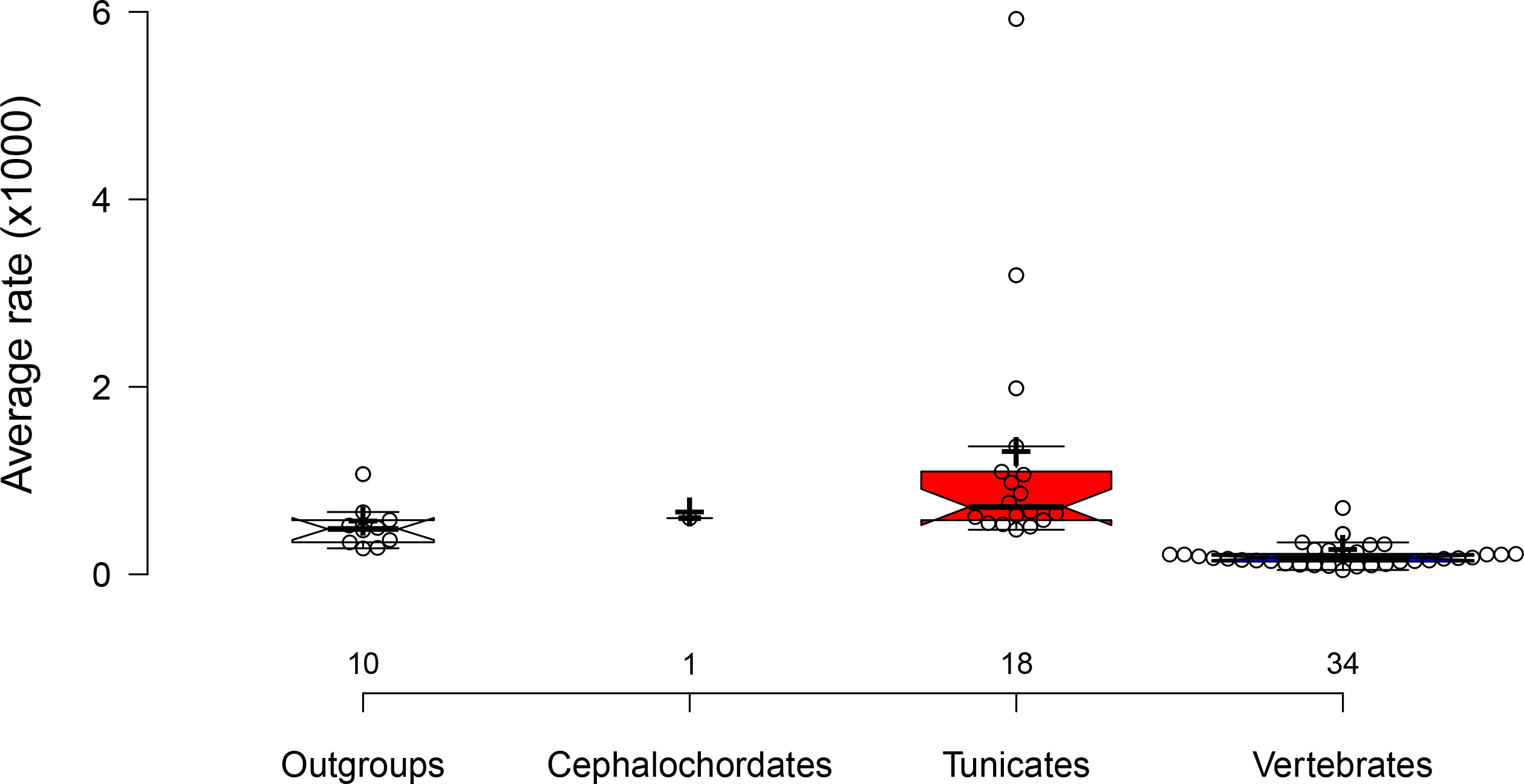
Evolutionary rate variation across sampled species. The bar plots represent average rate estimates (in number of substitutions per site per million years) obtained for the 63 terminal taxa regrouped by taxonomy. The rates were calculated using a rate-autocorrelated log-normal (LN) relaxed molecular clock model under the CAT-GTR+Γ_4_ mixture model with a birth-death prior on the diversification process and 13 soft calibration constraints. Data points are plotted as open circles with n = 10, 1, 18, 34 sample points in each taxonomic categories. Centre lines show the medians, crosses represent sample means, and box limits indicate the 25th and 75th percentiles with whiskers extending 1.5 times the interquartile range from the 25th and 75th percentiles. The width of the boxes is proportional to the square root of the sample size. This figure was made with BoxPlotR [81].

Even though the difference in fit between the two relaxed clock models was not significant for our dataset, in general LN provided more consistent dating estimates than UGAM with respect to the mean divergence dates of numerous vertebrate groups reported in the latest phylogenomic study of jawed vertebrates [35]. Notably, as observed in a previous phylogenomic study of tetrapods [74], the application of the UGAM relaxed clock model provided unrealistically recent estimates with respect to the maximum node age for the origin of turtles (LN mean age + SD: 180 ± 19 Mya [95% credibility interval: 220-146]; UGAM: 59 ± 41 Mya [173 - 16]) (Table 1; Fig. 3 and Additional file 2: Figure S1). The UGAM model also tended to systematically provide much wider 95% credibility intervals than LN with several of them actually spanning hundreds of millions of years (Table 1; Fig. 1; Additional file 2: Figure S1). Given the uncertainty associated with the dating results obtained using the UGAM model of clock relaxation, we focused our discussion below on results obtained with the more robust autocorrelated LN model, which we considered as our currently most reliable dating estimates.

**Figure 3.**
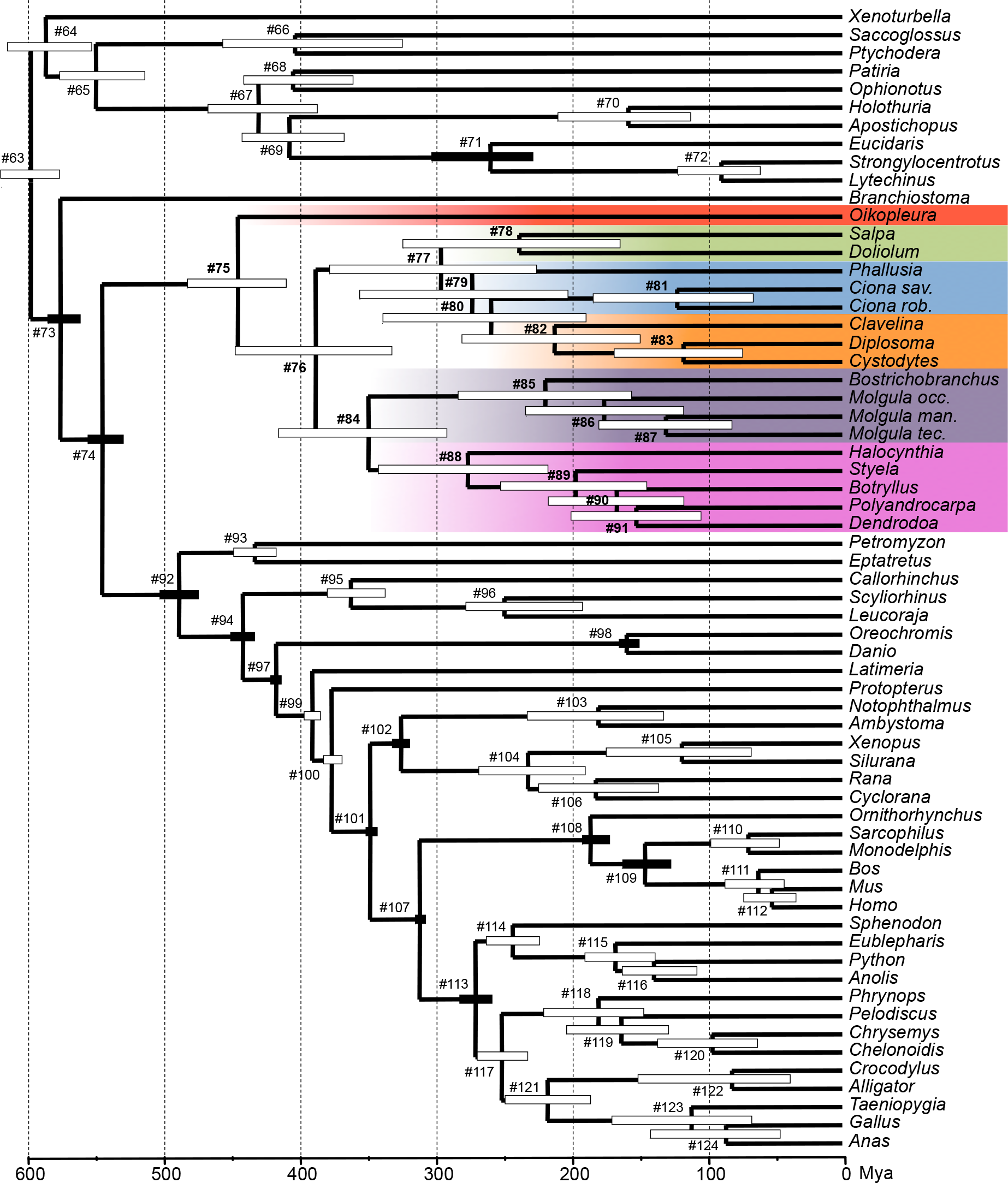
A molecular timescale for tunicates within chordates. The Bayesian chronogram has been obtained using a rate-autocorrelated log-normal (LN) relaxed molecular clock model using PhyloBayes under the CAT-GTR+Γ_4_ mixture model, with a birth-death prior on the diversification process, and 13 soft calibration constraints. Node bars indicate the uncertainty around mean age estimates based on 95% credibility intervals. Plain node bars indicated nodes used as a priori calibration constraints. Numbers at nodes refer Table 1.

**Table 1.** Molecular estimates of divergence dates (in Mya). The reported values represents mean divergence dates and associated standard deviations and 95% credibility intervals obtained from a Bayesian relaxed molecular clock under the LN and UGAM models coupled with a CAT-GTR+Γ_4_ mixture model.

| Nodes | LN<br>CAT-GTR+$\Gamma_4$ | | UGAM<br>CAT-GTR+$\Gamma_4$ | |
| --- | --- | --- | --- | --- |
| | Mean ± SD | 95% CredI | Mean ± SD | 95% CredI |
| #63 Deuterostomia | 599 ± 11 | [621 - 579] | 671 ± 108 | [985 - 576] |
| #64 Xenambulacraria | 588 ± 16 | [616 - 555] | 600 ± 89 | [849 - 467] |
| #65 Ambulacraria | 551 ± 16 | [578 - 516] | 517 ± 72 | [677 - 403] |
| #66 Hemichordata | 404 ± 34 | [458 - 326] | 206 ± 101 | [427 - 63] |
| #67 Echinodermata | 431 ± 21 | [469 - 388] | 403 ± 47 | [507 - 323] |
| #68 | 406 ± 21 | [442 - 363] | 284 ± 83 | [433 - 121] |
| #69 | 408 ± 20 | [443 - 368] | 360 ± 42 | [450 - 287] |
| #70 | 158 ± 22 | [210 - 112] | 117 ± 51 | [249 - 42] |
| #71 Echinoidea* | 260 ± 18 | [303 - 229] | 266 ± 28 | [342 - 222] |
| #72 | 89 ± 15 | [121 - 61] | 85 ± 49 | [195 - 20] |
| #73 Chordata* | 578 ± 6 | [586 - 563] | 575 ± 7 | [586 - 558] |
| #74 Olfactores* | 547 ± 6 | [557 - 532] | 545 ± 11 | [564 - 523] |
| **#75 Tunicata** | **447 ± 20** | **[484 - 411]** | **450 ± 26** | **[495 - 398]** |
| **#76** | **389 ± 32** | **[449 - 333]** | **388 ± 30** | **[439 - 326]** |
| **#77** | **296 ± 44** | **[379 - 226]** | **311 ± 40** | **[380 - 228]** |
| **#78 Thaliacea** | **238 ± 44** | **[324 - 164]** | **218 ± 54** | **[318 - 118]** |
| **#79** | **274 ± 44** | **[356 - 203]** | **272 ± 43** | **[351 - 176]** |
| **#80** | **259 ± 43** | **[340 - 190]** | **246 ± 44** | **[330 - 154]** |
| **#81 Ciona** | **122 ± 33** | **[184 - 65]** | **97 ± 44** | **[196 - 32]** |
| **#82 Aplousobranchia** | **212 ± 39** | **[281 - 150]** | **196 ± 44** | **[282 - 120]** |
| **#83 Cystodytes / Clavelina** | **117 ± 27** | **[168 - 73]** | **121 ± 32** | **[189 - 66]** |
| **#84 Stolidobranchia** | **350 ± 36** | **[416 - 292]** | **326 ± 39** | **[396 - 245]** |
| **#85 Mogulidae** | **219 ± 35** | **[285 - 156]** | **203 ± 48** | **[297 - 122]** |
| **#86 Molgula** | **176 ± 32** | **[233 - 118]** | **145 ± 40** | **[239 - 82]** |
| **#87 M. manhattensis / M. tectiformis** | **130 ± 26** | **[179 - 82]** | **94 ± 31** | **[162 - 44]** |
| **#88 Styelidae+Pyuridae** | **277 ± 35** | **[343 - 218]** | **228 ± 49** | **[323 - 139]** |
| **#89** | **197 ± 28** | **[252 - 145]** | **152 ± 41** | **[249 - 84]** |
| **#90** | **167 ± 25** | **[217 - 118]** | **113 ± 34** | **[187 - 59]** |
| **#91 Polyandrocarpa / Dendrodoa** | **152 ± 24** | **[200 - 105]** | **84 ± 30** | **[156 - 37]** |
| #92 Vertebrata | 490 ± 7 | [504 - 476] | 481 ± 13 | [510 - 460] |
| #93 Cyclostomata | 434 ± 8 | [449 - 418] | 277 ± 94 | [430 - 101] |
| #94 Gnathostomata* | 443 ± 4 | [452 - 435] | 437 ± 9 | [459 - 424] |
| #95 Chondrichthyes | 363 ± 11 | [380 - 338] | 192 ± 96 | [394 - 62] |
| #96 | 249 ± 22 | [277 - 192] | 88 ± 59 | [261 - 23] |
| #97 Osteichthyes* | 418 ± 2 | [422 - 416] | 419 ± 2 | [422 - 416] |
| #98 Clupeocephala* | 159 ± 4 | [165 - 150] | 157 ± 5 | [165 - 150] |
| #99 | 391 ± 3 | [397 - 386] | 393 ± 15 | [415 - 360] |
| #100 | 377 ± 3 | [383 - 371] | 374 ± 16 | [405 - 346] |
| #101 Tetrapoda* | 349 ± 2 | [351 - 345] | 341 ± 6 | [350 - 330] |
| #102 Amphibia | 326 ± 3 | [332 - 320] | 246 ± 30 | [299 - 200] |
| #103 | 180 ± 26 | [232 - 132] | 71 ± 51 | [190 - 14] |
| #104 Batrachia* | 232 ± 21 | [268 - 190] | 123 ± 48 | [225 - 47] |
| #105 | 118 ± 29 | [174 - 68] | 42 ± 32 | [132 - 9] |
| #106 | 182 ± 25 | [224 - 136] | 71 ± 37 | [160 - 21] |
| #107 Amniota* | 312 ± 1 | [315 - 310] | 319 ± 5 | [329 - 312] |
| #108 Mammalia* | 186 ± 5 | [192 - 172] | 176 ± 8 | [191 - 163] |
| #109 Theria* | 146 ± 9 | [163 - 127] | 143 ± 14 | [170 - 123] |
| #110 | 69 ± 13 | [96 - 47] | 47 ± 35 | [128 - 8] |
| #111 | 62 ± 11 | [86 - 43] | 55 ± 31 | [127 - 15] |
| #112 | 52 ± 10 | [73 - 35] | 35 ± 24 | [100 - 9] |
| #113 Diapsida* | 271 ± 6 | [282 - 259] | 278 ± 14 | [300 - 256] |
| #114 Lepidosauria | 243 ± 10 | [261 - 224] | 166 ± 69 | [283 - 57] |
| #115 | 168 ± 14 | [189 - 138] | 88 ± 47 | [203 - 28] |
| #116 | 139 ± 14 | [162 - 107] | 55 ± 35 | [152 - 16] |
| #117 | 252 ± 9 | [269 - 233] | 154 ± 69 | [280 - 55] |
| #118 Testudines | 180 ± 19 | [220 - 146] | 59 ± 41 | [173 - 16] |
| #119 | 163 ± 19 | [204 - 128] | 38 ± 28 | [120 - 11] |
| #120 | 96 ± 18 | [136 - 62] | 15 ± 14 | [54 - 4] |
| #121 Archosauria | 218 ± 16 | [249 - 186] | 102 ± 51 | [238 - 37] |
| #122 | 81 ± 29 | [150 - 38] | 23 ± 22 | [85 - 4] |
| #123 Aves | 111 ± 27 | [170 - 67] | 45 ± 28 | [120 - 14] |
| #124 Crocodylia | 86 ± 24 | [142 - 46] | 23 ± 18 | [75 - 6] |

\*Calibration constraints

### A tentative timescale for tunicate evolution within chordates

The Bayesian chronogram obtained using the LN relaxed molecular clock model and the site-heterogeneous CAT-GTR+Γ_4_ mixture model of amino acid sequence evolution is presented in Figure 3. This phylogenomic timescale showed that major tunicate clades appeared early in chordate evolutionary history. The earliest split between appendicularians and all other tunicates was dated back to ca. 450 Mya (mean age + SD: 447 ± 20 Mya [95% credibility interval: 484 - 411]), followed by the divergence between stolidobranchs and the clade grouping thaliaceans + phlebobranchs + aplousobranchs ca. 390 Mya (389 ± 32 Mya [449 - 333]), and the separation of stolidobranchs into Molgulidae and Styelidae + Pyuridae ca. 350 Mya (350 ± 36 Mya [416 - 292]) (Table 1; Fig. 3). Even more recent divergences such as the ones between congeneric species within *Ciona* and *Molgula* occurred more than 100 Mya.

Given the relative uncertainty on the phylogenetic position of *Xenoturbella,* complementary LN relaxed molecular clock analyses were also conducted using *Xenoturbella* as an outgroup. As the dating results previously obtained with the CAT-F81+Γ_4_ and CAT-GTR+Γ_4_ models with the original rooting were extremely similar (linear regression on mean dates: R^2^ = 0.99), we performed these additional analyses under the less computationally intensive CAT-F81+Γ_4_ model. With the new rooting configuration, the inferred mean divergence dates between the two alternative rooting schemes were globally highly correlated within Chordates (linear regression: R^2^ = 0.89). An almost exact correspondence was found for Vertebrates that contain most of the calibration points (linear regression: R^2^ = 1.00). For Tunicates, within which there is unfortunately no available calibration constraint, the correlation remained very strong (linear regression: R^2^ = 0.95). The divergence dates within tunicates were on average older with the *Xenoturbella* rooting, while they remained in their vast majority within the original 95% credibility intervals (Additional file 3: Figure S2). An alternative rooting by *Xenoturbella* thus does not affect our main conclusions that divergence dates among the major tunicate lineages are ancient.

Our estimated divergence dates in tunicates were nevertheless associated with fairly large 95% credibility intervals, probably because of the lack of internal fossil calibrations within tunicates, in contrast to the well-calibrated vertebrates. It has recently been pointed that given the uncertainty associated with molecular dating estimates, building evolutionary narratives would be premature for early animal evolution [75]. In our case, we argue that in the absence of a trustable tunicate fossil record [33], our tentative molecular timescale constitutes the first and only currently available approach to provide a much-needed relative comparison of divergence times between the major lineages of tunicates and vertebrates. Such a comparison is subject to considerable uncertainty but it has nevertheless revealed several deep divergences occurring at comparable geological times between the two groups (Fig. 3; Table 2). For instance, between tunicates (*Ciona* / *Oikopleura*) and gnathostomes (*Homo* / *Callorhinchus*) around 450 Mya; thaliaceans (*Salpa* / *Doliolum*) and lepidosaurs (*Sphenodon* / *Anolis*) around 240 Mya; and between stolidobranchs (*Molgula* / *Botryllus*) and tetrapods (*Homo* / *Xenopus*) around 350 Mya.

**Table 2.**
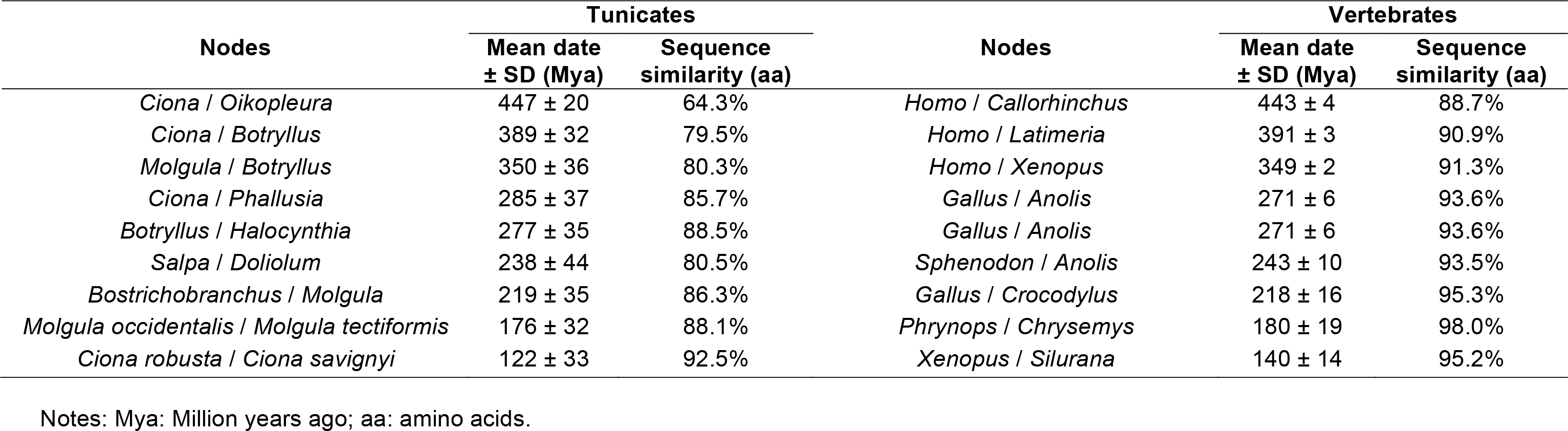
Parallel divergences between model tunicates and vertebrates. The reported values indicate mean divergence dates and associated standard deviations obtained from a Bayesian relaxed molecular clock under the CAT-GTR+G_4_ and the percentage of amino acid sequence identity for each couple.

The relatively ancient origins of the different tunicate lineages revealed by our molecular dating estimates have two broader implications. First, there seems to be a larger gap than previously thought between tunicate and vertebrate taxonomic ranks which exacerbates the inadequacy of their direct comparison. For example, when a vertebrate genus usually spans less than 40 million years [76], a tunicate genus (e.g. *Molgula*) can span up to two hundred million years (Fig. 3; Table 1). The meaning of Linnean categorical ranks and their temporal inconsistencies among clades have been largely discussed [76], as recently illustrated by the debate around the taxonomic status of the main Chordate lineages [77–79]. The parallel we draw here between tunicates and vertebrates should nevertheless help tunicate developmental biologists to interpret their results in light of the large divergences that might exist between tunicate model species despite their classification in the same genus. Second, the ancient age of their major divergence events can heavily complicate orthology assessment among tunicates, as well as between tunicates and vertebrates, thus reducing the quality of genome annotations. Indeed, the fast-paced molecular evolution of tunicates prevents the identification of some genes by simple similarity methods (e.g. BLAST), even when orthologs do exist in databases. For instance, in terms of evolutionary depth, a comparative study of the genus *Molgula* is roughly equivalent to a comparative study among turtles representing about 180 Myr of evolution. In terms of amino acid sequence divergence, the differences are much more pronounced between *Molgula occidentalis* / *Molgula tectiformis* (88.1% similarity) than between *Phrynops* / *Chrysemys* (98.0% similarity; Fig. 3, Table 1).

From an evo-devo perspective, the phylogenetic framework and tentative timescale presented here lead to an apparent paradox. Like most nematodes [80], the embryos of each ascidian species develop in a stereotyped manner, based on the use of invariant cell lineages [7]. Unlike nematodes however, ascidian stereotyped cell lineages are shared between evolutionarily distant species such as *Ciona robusta* (Enterogona) and *Halocynthia roretzi* (Pleurogona) [11]. The extreme morphological conservation of ascidian embryogenesis therefore contrasts with the high rates of protein divergence observed in their genomes. This paradox raises questions about the underlying mechanisms involved in developmental regulation of these animals with highly dynamic genomes. In this context, our reference phylogenetic tree and divergence date estimates among tunicate lineages could be used as an evolutionary framework to select model species sufficiently close to one another (i.e. retaining sufficient phylogenetic information) for future comparative genomic analyses assessing orthology by gene tree reconciliation and estimating evolutionary rate variations among gene ontology categories.

### Conclusion

This study represents the first large-scale phylogenomic analysis including all major tunicate lineages based on transcriptomic data. The resulting phylogenetic framework and tentative timescale constitute a necessary first step towards a better understanding of tunicate systematics, genomics, and development, and in a broader context, of chordate evolution and developmental biology.

## Declarations

### Ethics approval and consent to participate

Not applicable.

### Consent for publication

All authors agree with the publication of this article.

### Availability of data and material

All data generated or analysed during this study are included in this published article and its supplementary information files. Raw sequencing reads have been deposited under NCBI Bioproject PRJNA414754. Transcriptome assemblies, alignments, and trees are publicly available from a Github repository (https://github.com/psimion/SuppData_Delsuc_BMCBiol_2018_Dating_Tunicata).

### Authors’ contributions

F.D., H.P., and E.J.P.D. designed the study. F.D., G.T., X.T., S.L.-L., and J.P. collected and prepared biological material. F.D., G.T., M.-K.T., J.P., P.L., and E.J.P.D. organized sequence data collection. H.P. and G.T. constructed the supermatrix. F.D., H.P., P.S., and E.J.P.D analysed data. F.D., H.P., P.S., and E.J.P.D. drafted the manuscript. All authors contributed to the final version of the manuscript and gave final approval for publication.

## Acknowledgments

We thank Nicolas Galtier for giving early access to *Clavelina* and *Cystodytes* Illumina data. We are thankful to Stefano Tiozzo, Hector Escriba, Sébastien Darras, Frédérique Viard, and the diving staffs of the marine stations of Villefranche sur Mer, Banyuls, and Roscoff for their help in sample collection. We also thank two anonymous referees for their comments. This is contribution ISEM 2018-024 of the Institut des Sciences de l’Evolution de Montpellier.

---

**Figure S1.** Bayesian chronogram obtained using an uncorrelated gamma (UGAM) relaxed molecular clock model using PhyloBayes under the CAT-GTR+Γ_4_ mixture model, with a birth-death prior on the diversification process, and 13 soft calibration constraints. Node bars indicate the uncertainty around mean age estimates based on 95% credibility intervals. Plain black node bars indicated nodes used as a priori calibration constraints. Numbers at nodes refer to Table 1.

**Figure S2.** Bayesian chronogram obtained using an autocorrelated lognormal (LN) relaxed molecular clock model using PhyloBayes under the CAT-F81+Γ_4_ mixture model, with a birth-death prior on the diversification process, 13 soft calibration constraints, and an alternative rooting by *Xenoturbella*. Node bars indicate the uncertainty around mean age estimates based on 95% credibility intervals. Plain black node bars indicated nodes used as a priori calibration constraints. Numbers at nodes refer to Table 1.

